# Cellular senescence and quiescence are associated with altered ribosomal RNA methylation and processing

**DOI:** 10.1101/2020.04.01.019653

**Authors:** Guohuan Yang, Clemens Heisenberger, Isabelle C. Kos-Braun, Norbert Polacek, Johannes Grillari, Markus Schosserer, Martin Koš

## Abstract

The 2’-O-methylation (2’-O-Me) of ribosomal RNA (rRNA) shows plasticity potentially associated with specific cell phenotypes. We used RiboMeth-seq profiling to reveal 2’-O-Me patterns specific to stress induced premature senescent (SIPS), quiescent and proliferating human dermal fibroblasts. The altered methylation levels in SIPS and quiescence partially correlated with the expression of specific snoRNAs but not fibrillarin. Senescence and quiescence were accompanied by a trend towards preferred usage of one of the two alternative ribosome biogenesis pathways.

## Introduction

The accumulation of senescent cells is one of the main drivers of biological ageing 31 mainly due to the characteristic pro-inflammatory senescence-associated secretory phenotype (SASP) (Coppe et al., 2010). Eliminating senescent cells extends the healthy lifespan of mice (Baker et al., 2016; Xu et al., 2018). Thus, strategies to deplete senescent cells or block the SASP offer therapeutic opportunities. Ribosomes represent a promising novel target, as delayed rRNA processing promotes induction of senescence by activation of p53 (Nishimura et al., 2015), and accumulation of pre-ribosomes maintains the senescent state by engaging retinoblastoma protein (Lessard et al., 2018). The protein composition of ribosomes in senescent cells was found to vary (Shi et al., 2017), but other ribosome features, such as rRNA modifications, were not yet explored.

The 2’-O-methylation (2’-O-Me), the most abundant modification in rRNA, is introduced by ribonucleoprotein complexes of four conserved proteins, including the methyltransferase fibrillarin (FBL), and a small nucleolar RNA (snoRNA) specifying the methylated site (Kiss, 2002). Recently, the plasticity of the 2’-O-Me modification of rRNA in response to stress was shown to contribute to the heterogeneity of ribosomes and affect their translational activity (Erales et al., 2017; Krogh et al., 2016). Here we show that the rRNA 2’-O-methylation is altered in senescent and quiescent human cells.

## Results and discussion

We induced senescence in the human dermal fibroblasts (HDF) from three different donors by exposure to hydrogen peroxide (Lämmermann et al., 2018) and confirmed their senescent phenotype by changes in morphology, increased senescence-associated β-Galactosidase activity (SA-β-gal) and absence of proliferation by incorporation of 5-Bromo-2’-deoxyuridine (BrdU) (Supp. Inf. Fig. S1). The contact-inhibited quiescent (Q) and proliferating (P) cells derived from each donor served as controls.

We used RiboMeth-seq to profile 104 2’-O-Me sites in the rRNA of SIPS, Q and P cells and assigned the RMS score (0 = non-methylated, 1 = fully methylated) to each modified position as described (Krogh et al., 2016) (Fig. 1 and Supp. Inf. Tables S2, S3). While the majority of sites were highly methylated (RMS > 0.9), the scores of nine positions in SIPS and Q cells appeared lower (SSU-C1391, LSU-C3680) or higher (SSU: C797, G867, C1272 and G1447; LSU: G1303, G3723 and G4588) compared to P cells (Fig. 1). The methylation at SSU-C797, G867, C1272 and LSU-G1303 was nearly identical in Q and SIPS but different in P cells, pointing to a general role in proliferation. The SSU-C1391, G1447, and LSU-C3680, G3723 methylation in SIPS differed from P or Q cells suggesting a specific association with SIPS, however the biological variance between donors at these sites precludes a conclusion from the current data (Fig. 1; Supp. Inf. Tables S2, S3).

**Fig. 1:**
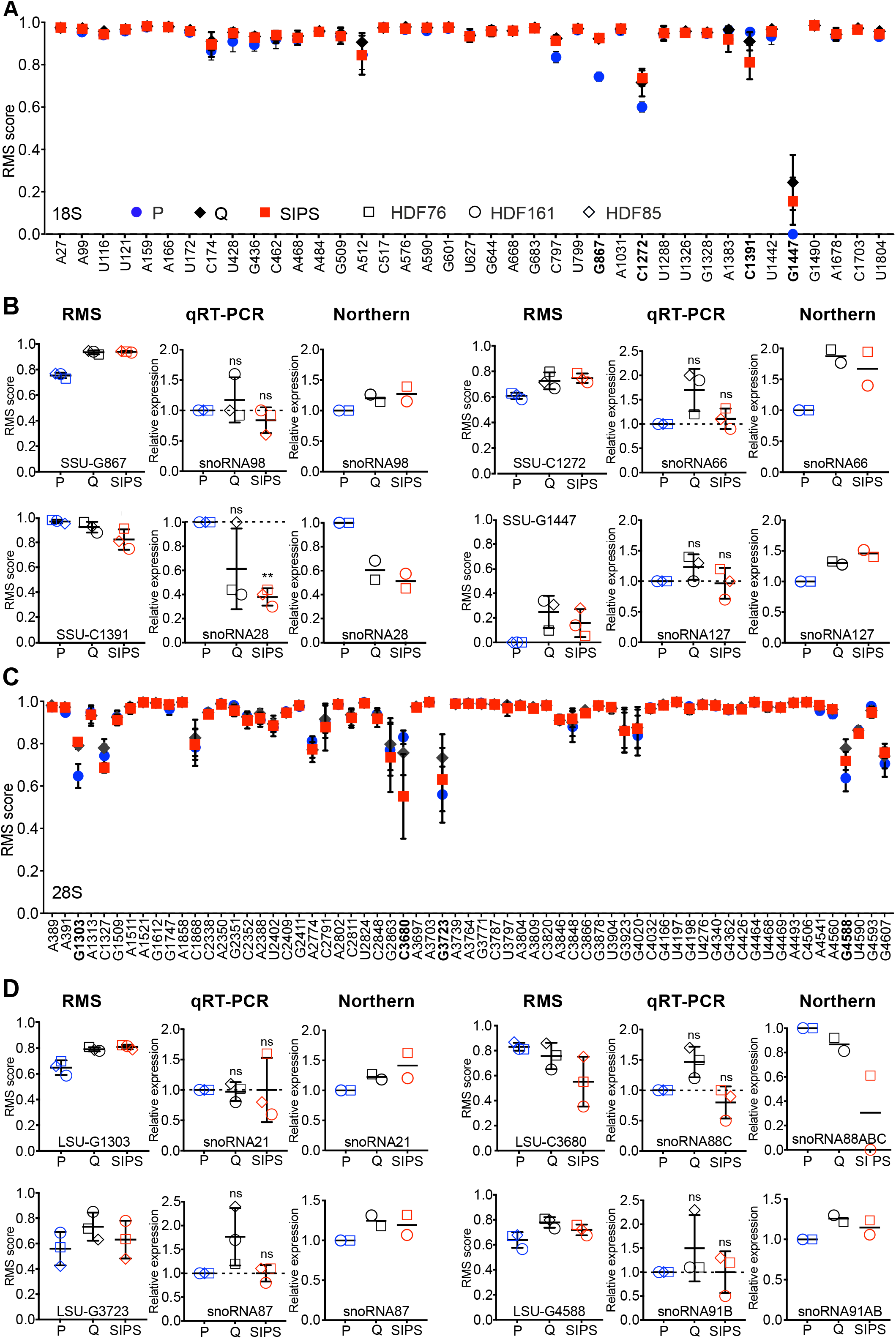
RiboMeth-seq and snoRNA expression. **(A, C)** Average RMS scores for each 2’-O-methylated nucleotide in 18S (A) and 28S (C) rRNAs from proliferating (P), quiescent (Q) and senescent (SIPS) HDFs from three donors (HDF76, HDF161, HDF85). Error bars represent standard deviation. **(B and D)** Relative levels of snoRNAs guiding modification at the variable sites. Values were corrected for loading by snoRNA57 (northern) and 5.8S RNA (qRT-PCR).

The RMS scores were overall slightly higher in Q and SIPS than P cells. In a recent study, the expression levels of FBL affected methylation at specific positions (Sharma et al., 2017). However, in our model the methylation in Q/SIPS and P cells did not correlate with FBL expression, which did neither change at mRNA nor at protein level in in any of the tested conditions (Supp. Inf. Fig. S1D-F).

In contrast, methylation at the hypomodified sites correlated with expression of snoRNAs guiding modification at those sites. Even small methylation changes were accompanied by comparable alterations in the snoRNAs levels (Fig. 1BD, Supp. Inf. Fig. S2, S3). Importantly, the expression of all snoRNAs for the hypomodified sites was very low and thus the hypomethylation might be due to the limiting levels of the snoRNAs.

Contrary to previous reports of delayed early rRNA processing in cellular senescence (Lessard et al., 2018; Nishimura et al., 2015), the 47 pre-rRNA did not accumulate in SIPS revealing that neither transcription nor early processing were affected (Fig. 2ABC). Conversely, the relative levels of 21S and 18S-E pre-rRNAs decreased in SIPS, indicating an altered kinetics of the late SSU biogenesis (Fig. 2DE). Intriguingly, Q and SIPS cells showed a trend towards preference for the rRNA processing pathway B, represented by an increase in the 30S:41S pre-rRNA ratio (41S and 30S pre-rRNAs are produced only by the pathway A or B respectively) (Fig. 2A and 2FGH). A similar switch between two pre-rRNA processing pathways in response to stress was described in yeast (Kos-Braun et al., 2017). Alternative pathways can provide means to produce distinct ribosomes.

**Fig. 2:**
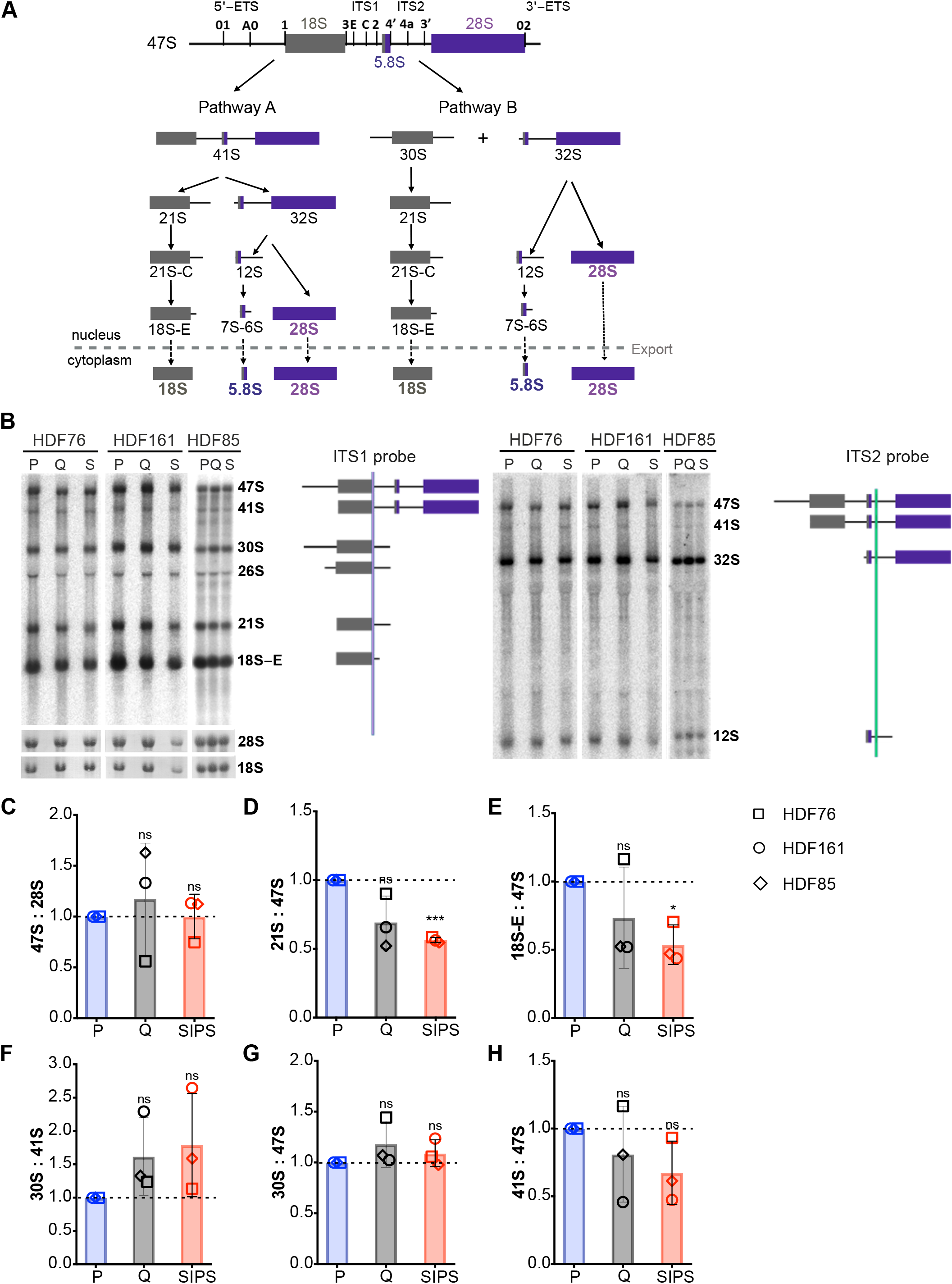
Pre-rRNA processing in SIPS. **(A)** Scheme of the human pre-rRNA processing (modified from (Mullineux & Lafontaine, 2012). **(B)** Northern blots of total RNA using two probes to detect different pre-rRNAs shown on the right. (**C-H)** Quantification of changes in pre-rRNAs expressed as ratios: **(C)** 47S:28S; **(D)** 18S-E:47S; **(E)** 21S:47S; **(F)** 30S:41S; **(G)** 30S:47S; **(H)** 41S:47S.

In summary, we identified nine 2’-O-Me sites that appear to be differentially modified in non-proliferating cells (Q and SIPS). Their methylation correlated with the snoRNAs levels that are likely at limiting concentrations. Upon growth arrest, the pre-rRNA processing in SIPS and Q cells shifted towards the pathway B. Our results provide a basis for further study of ribosomes in cellular senescence as plausible targets for novel therapeutic interventions.

## ACKNOWLEDGEMENTS

We thank Uschi Göbels and Elena Stelzer for technical assistance. GY was supported by the China Scholarship Council (201608310107). The work was supported by Deutsche Forschungsgemeinschaft 280594475 to MK, Austrian Science Fund (FWF) I2514 to JG and the Swiss National Science Foundation 310030E-162559/1 to NP; and by FWF and Herzfelder’sche Familienstiftung P30623-B26 to MS.

## AUTHOR CONTRIBUTIONS

GY, CH and IKB performed experiments and analyzed data. NP, JG, MS and MK designed experiments and analyzed data. GY, MS and MK wrote the paper.

