## Supplementary Material for "Cellular senescence and quiescence are associated with altered ribosomal RNA methylation and processing"

### SUPPORTING INFORMATION

- **Experimental procedures**
- **Supplementary Figures (S1-S3)**
- **Supplementary Tables (S1-S5)**

### EXPERIMENTAL PROCEDURES

#### *Cell culture*

Primary human dermal fibroblasts (HDF) from three healthy donors (HDF76, HDF85 and HDF161) were obtained from Evercyte GmbH (Vienna, Austria). Details of all donors are listed in Supporting Information Table S1. Cells were cultured with DMEM/Ham's F-12 (1:1 mixture) (F4815, Biochrom) supplemented with 10% fetal calf serum (F7524, Sigma) and 4 mM L-glutamine (G7513; Sigma) under ambient oxygen, 7% $CO_2$ and 37°C. Cell counting was done with an automated cell counter (Vi-CELL XR, Beckman Coulter).

Proliferating cells were passaged twice a week and were detached by incubation with 0.1% trypsin and 0.02% EDTA at 37°C for 5 min and split at ratios between 1:2 or 1:3 depending on cell type and growth rate.

For quiescent/SIPS state, cells were seeded at a density of 3500 cells/cm². Quiescent cells were then left to grow until contact inhibited (while exchanging medium once a week) in parallel to SIPS cells being treated to induce senescence (see below).

#### *Stress-induced premature senescence (SIPS) by hydrogen peroxide exposure or doxorubicin treatment*

Following the protocol established in our lab (Lämmermann et al. 2018) we prepared SIPS cells from three donors. Due to differences in their replicative lifespan (Supporting Information Table S1), cells of different donors at corresponding PD were seeded at a density of 3500 cells/cm² and left to attach to the surface overnight. The following day, stress treatment was started. 30% $H_2O_2$ stock solution (Sigma) was freshly diluted and added to the medium to a final concentration of 60-80 μM. After one hour incubation at 37°C, the medium was replaced to remove residual $H_2O_2$. This hydrogen peroxide treatment was performed on four consecutive days, followed by two days break, another five consecutive days of treatment, and finally 5-6 days recovery. SIPS by hydrogen peroxide exposure was used for all assays except the quantification of snoRNA expression by qPCR.

Doxorubicin medium (Doxo-Medium) was freshly prepared with a final concentration of 150 nM to 200 nM depending on the donor. Seeding was performed as for SIPS-induction with $H_2O_2$. On the following day, the medium was replaced with Doxo-Medium and incubated at 37°C, 7% $CO_2$ for 4 days. Thereafter, the medium was replaced with fresh Doxo-Medium and cells were incubated for another seven consecutive days. After doxorubicin exposure, cells were supplied with fresh regular growth medium once a week to recover for 3 weeks. Total RNA from these cells was used to quantify snoRNA expression by qPCR.

Induction of premature senescence was confirmed by the altered cellular morphology and senescence-associated β-galactosidase (SA-β-gal) activity. The cell cycle arrest was confirmed by BrdU-incorporation.

***SA-β-gal staining***

SA-β-gal staining was performed as previously described (Dimri et al. 1995). 10x images were captured for each well at ×100 magnification and positive and negative cells of P and SIPS cells were counted after randomization. 15 individual frames with at least 300 cells total were counted per condition. Counting was performed twice and averaged.

#### *5-bromo-2'-deoxyuridine (BrdU) labeling*

Cells were incubated in DMEM/Ham's supplemented with 10 µM BrdU (Bromodeoxyuridine) (Sigma Aldrich) for 24 hours. After harvesting with trypsin, cells were fixed using ice cold 70% ethanol for at least one hour at 4°C. Denaturation of the DNA for 30 min with 2 M HCl and 1% Triton X-100 (Sigma Aldrich) was performed, followed by neutralization with 0.1 M Na-Borat, pH 8.5. Pellets were resuspended in TBS (0.5% Tween20, 1% BSA in 1 x PBS) containing anti-BrdU antibody 1:50 (BD Biosciences, USA, 347580) and incubated for 30 min at room temperature. After washing with TBS and staining with anti-mouse FITC-conjugated secondary antibody (Sigma F-8264) for 30 minutes, the pellet was washed with TBS and resuspended in 1x PBS containing 2.5 µg/ml propidium iodide (PI) (Sigma Aldrich). For compensation, cells were stained with either PI or BrdU alone. Analysis was performed by flow cytometry (Gallios Beckman Coulter).

#### *RNA extraction and purification*

Cells were harvested and total RNA was isolated using TRIzol Reagent (Sigma) following the manufacturer's protocol. Further purification of total RNA was achieved by washing the RNA pellet with 70% ethanol twice. The RNA concentration was measured with an ND-1000 (NanoDrop) spectrometer.

#### *qPCR*

cDNA was synthesized from 50 ng total RNA using High-Capacity cDNA Reverse Transcription Kit (Life Technologies). Target gene expression levels were quantified from cDNA using 5x HOT FIREPol® EvaGreen® qPCR Mix Plus (Medibena) on a Rotor-Gene Q cycler (Qiagen). The reference gene GAPD was used for normalization of Fibrillarin.

Identification and quantification of individual snoRNAs was conducted with specific stem-loop reverse transcription primers, gene-specific forward primers and an universal reverse primer (Kramer 2011). For snoRNA quantification by qPCR, SIPS was induced by doxorubicin exposure. snoRNAs expression levels were analyzed by comparison to a standard curve and normalization of absolute expression levels to 5.8S rRNA.

Sequences of qPCR primers are provided in Supplementary Table S5.

***Preparation of RNA fragments library and RiboMetseq***

5 μg total RNA were hydrolyzed in 50 mM bicarbonate buffer pH 9.2 and 10 mM $MgCl_2$ at 95°C for 5 min, then put on ice. RNA cleavage was stopped by addition of 2 μl of 0.5 M EDTA pH 8.0. The RNA was precipitated by addition of 1/10 volume of 3M sodium acetate and 2 volumes of 100% ice-cold EtOH, washed with 70% ice-cold EtOH and resuspended in nuclease-free water. RNA fragments were separated on a denaturing 6% acryl:bis (19:1), 8M urea gel. The fragments in the size range of 20 to 40 nt were extracted from the gel. The 5´ ends of the RNA fragments were dephosphorylated with Shrimp Alkaline Phosphatase (rSAP) (New England Biolabs, M0371S) as described in the manufacturer’s protocol. The sequencing library was prepared using NEB SmallRNA for Illumina kit and sequenced as single-end, in two technical replicas at the Deep Sequencing Core Facility (BioQuant) of Heidelberg University. The RNA sequences were trimmed and aligned to the human genome (GRCh38) using Geneious software. The fraction of methylated rRNA and RMS scores were calculated as previously described (Birkedal et al. 2015).

***Northern blots and quantification***

For northern blotting, 4 μg of total RNA were separated on 1.2% glyoxal agarose gel or 8% polyacrylamide/urea gel. Separated RNA was hydrolyzed and then transferred to a Hybond+

nylon membrane (Roche) at 4°C using wet electro-transfer. The membrane was stained with methylene blue to confirm homogenous and efficient transfer. Probes were hybridized in Church hybridization buffer (0.36 M $Na_2HPO_4$, 0.14 M $NaH_2PO_4$, 1 mM EDTA, 7% SDS) at 38°C to 42°C overnight, washed and exposed to phosphor-imaging plates (Fuji). The imaging plates were scanned on Fuji FLA-5100 scanner and quantification was done using the AIDA software (Raytest), using 1D-quantification mode. Some snoRNA expression levels were low and required use of 2-3 different probes per snoRNA and several days of exposure. The signal of each snoRNA was normalized to snoRNA-57 (SSU-A99). snoRNA57 was chosen for normalization, because its signal was comparable to other snoRNAs and neither the RMS at its guided position nor the snoRNA expression changed significantly between the P, Q and SIPS cells. Results by normalization to 5.8S were similar. Northern blotting was performed with HDF76 and HDF161.

The scheme of pre-rRNA processing (Figure 2) was drawn based on a previous report (Mullineux & Lafontaine 2012).

***Western blots and quantification***

Harvested cells were lysed in RIPA-buffer (150 mM NaCl, 0.5% sodium deoxycholate, 1% NP-40, 0.1% SDS, 50 mM Tris/HCl pH 8.0). After sonification with a Bioruptor Plus sonicator (Diagenode) for 30 cycles (30s on/30s off), SDS- PAGE sample buffer (60 µM Tris/HCl pH 6.8, 2% SDS, 10% glycerol, 0.0125% bromophenol blue and 1.25% β -mercaptoethanol) was added and mix thoroughly. 4-15% Mini-PROTEAN® TGX Gels (BioRad) in Laemmli-Buffer (25 mM Tris, 250 mM glycine and 0.1% SDS) were used for electrophoresis. Transfer of separated protein bands to PVDF-membranes (Bio Rad) was performed at 25 V and 1.3 A for 3 min. Following blocking and antibody incubation, signal detection was done with Odyssey

Infrared Imager (LI-COR). Quantification of each band was done with ImageJ (Version 1.52 e) after brightness and contrast adjustments.

***Statistical Analysis***

GraphPad Prism 8.0 was used for all statistical analyses. The number of replicates (n = 3) for RiboMetSeq, qPCR for snoRNAs and fibrillarin expression, and northern blots for pre-rRNA processing refers to independent biological replicates representing three different donors (HDF76, HDF85 and HDF161). Results are presented as mean ± standard deviation. Statistical significance was tested with one sample *t*-test against an expected value of 1 for northern blots of rRNA precursors and RT-qPCR of snoRNAs, fibrillarin (FBL) mRNA expression and FBL protein expression; a two-tailed Student's *t*-test was used for SA-β-gal positive cells.

Throughout the text: $\alpha$=0.05, *$P < 0.05$, **$P < 0.01$, ***$P < 0.001$, ****$P < 0.0001$. Only statistically significant pairwise comparisons ($P < 0.05$) are indicated in all figures.

For northern blots of snoRNA expression, two biological replicates (HDF76 and HDF161) were performed. Values are presented as means.

***Data Availability***

The raw and processed sequencing data are available from the Gene Expression Omnibus database (https://www.ncbi.nlm.nih.gov/geo) under accession XXX. Reasonable requests for other data and reagents presented in this manuscript will be honored by the corresponding authors.

### SUPPLEMENTARY FIGURES

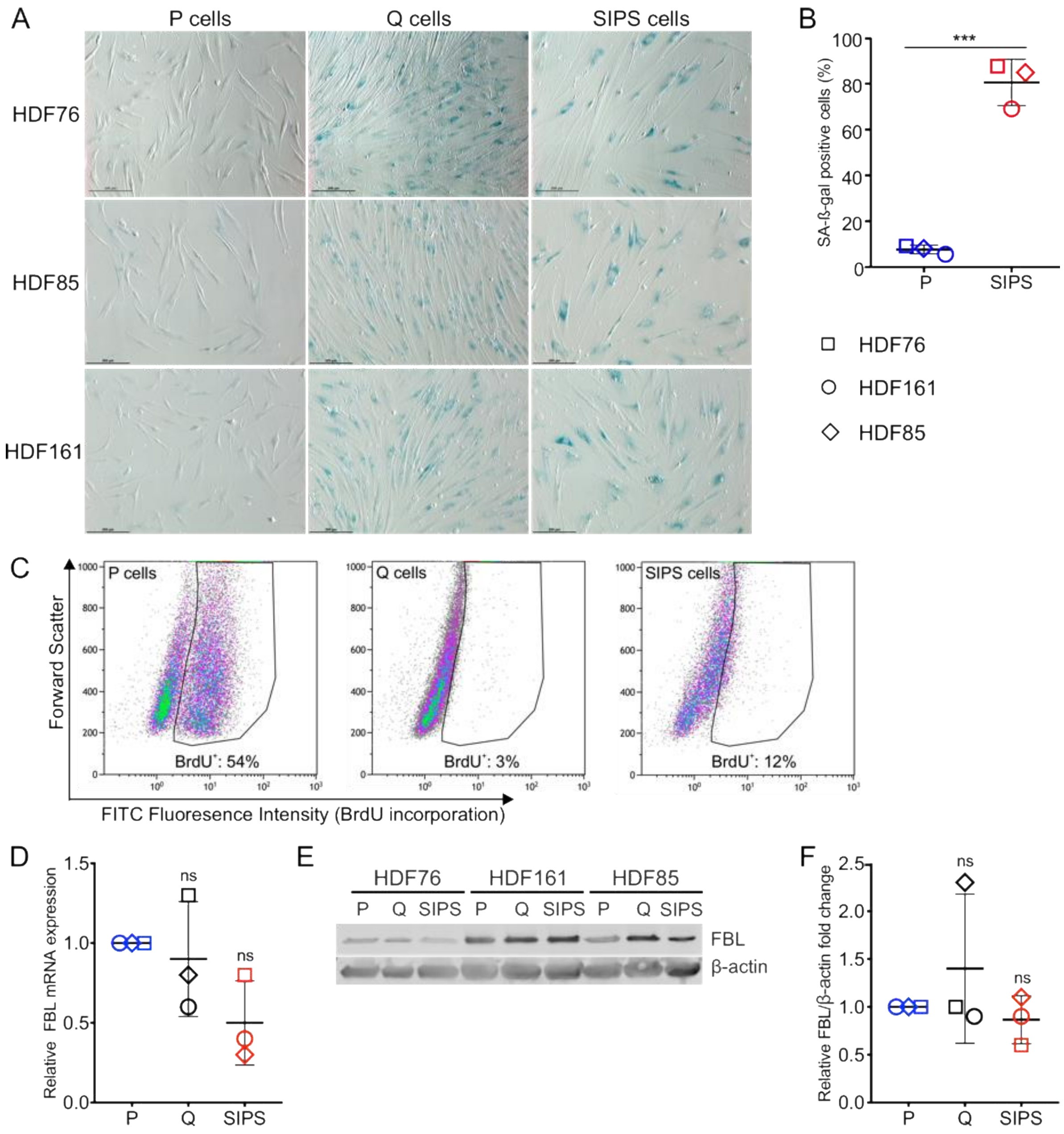


**Figure S1: Cellular senescence verification and Fibrillarin expression.** Senescence was confirmed by **(A)** and **(B)** senescence-associated β-galactosidase (SA-β-gal) activity. The scale bar for (A) represents 200 µm; (B) Quantification of SA-β-gal positive cells of P and SIPS cells for three donors (HDF76, HDF85, HDF161), Error bars represent mean ± standard deviation. A two-tailed Student's t-test was used to test for statistical significance. **(C)** BrdU-incorporation was measured by flow cytometry for all three donors with similar outcome.

HDF76 is shown as representative result. **(D)** FBL mRNA expression via RT-qPCR (n = 3 biological replicates) is shown. **(E)** Protein expression levels of FBL in P, Q and SIPS cells by western blot are shown, β-actin was used as loading control (n = 3 biological replicates). **(F)** Relative fibrillarin/β-actin fold change by quantification of western blot result from (E). One sample *t*-test against an expected value of 1 was done for (D) and (F). For all statistic test showing here, $\alpha=0.05$, ns = not significant, $^{*}P < 0.05$, $^{**}P < 0.01$, $^{***}P < 0.001$, $^{****}P < 0.0001$.

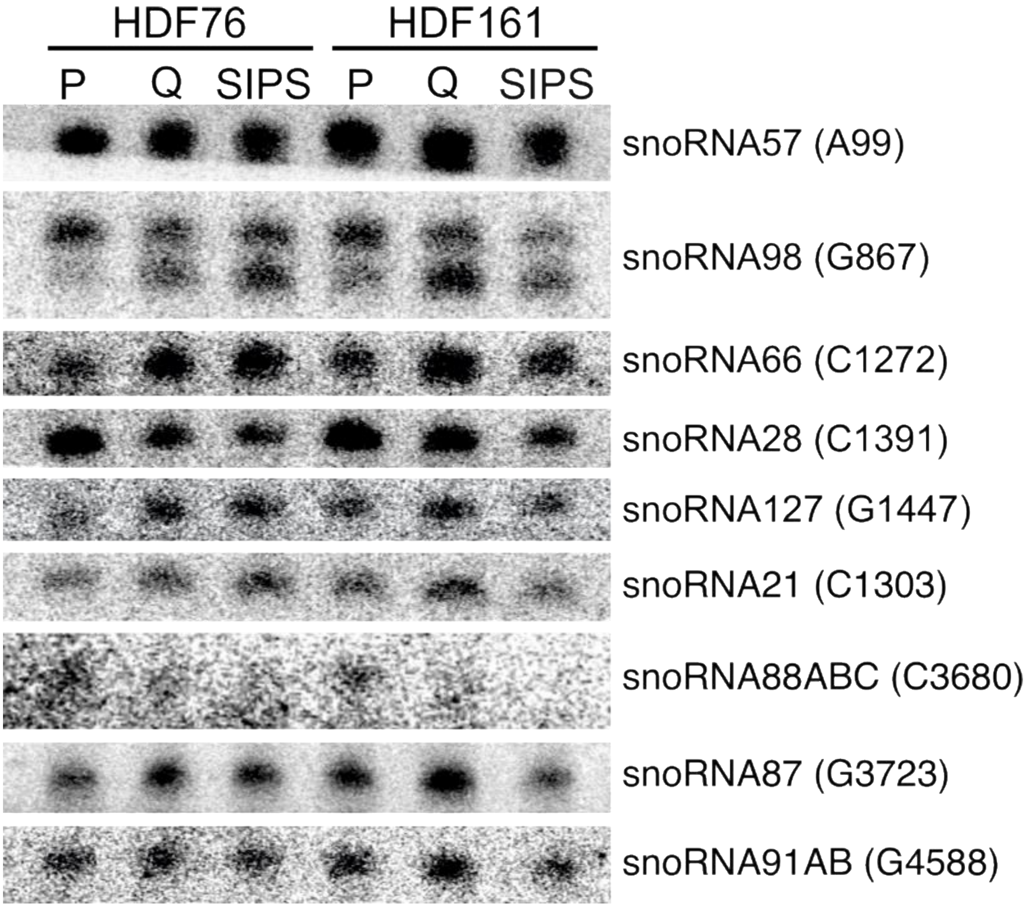


**Figure S2. Northern blot analysis of snoRNA expression.** Total RNA was separated on a 1.2% agarose-glyoxal gel, transferred to a nylon membrane and hybridized with probes complementary to snoRNAs.

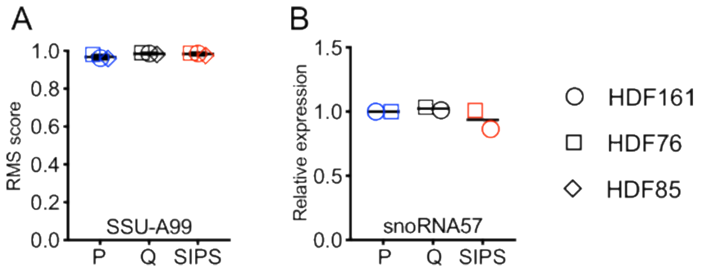


**Figure S3. Comparison of methylation and snoRNA expression for normalization control**. **(A)**The RMS scores at A99 are shown. **(B)** Quantification of snoRNA57 levels from the northern blot, the expression level in P cells was normalized to 1.

### SUPPLEMENTARY TABLES

#### TABLE S1: Characteristics of human dermal fibroblasts from three donors

| Name | Age of donor | Sex | Tissue origin | Max. PDs | Morphology of early-passage HDF | PD for SIPS |
|---|---|---|---|---|---|---|
| HDF161 | 65 y | Female | abdominoplasty | ~40* | reticular | 15 |
| HDF85 | 49 y | Female | abdominoplasty | ~35* | papillary | 13 |
| HDF76 | 58 y | Female | abdominoplasty | ~55* | papillary | 20.5 |

**TABLE S2: Average RMS scores at established 2'-O-Me sites in 18S rRNA from three donors.** Sites downregulated in SIPS cells in comparison with P/Q cells are highlighted in blue and italic and downregulated in P cells in comparison with Q/SIPS cells are highlighted in red and italic.

| | Average of HDF 76 & 161 & 85 | | | St. dev. HDF 76 & 161 & 85 | | |
|---|---|---|---|---|---|---|
| ***Established sites*** | **P** | **Q** | **SIPS** | **P** | **Q** | **SIPS** |
| **A27** | 0.987 | 0.988 | 0.989 | 0.011 | 0.010 | 0.009 |
| **A99** | 0.967 | 0.985 | 0.983 | 0.012 | 0.004 | 0.007 |
| **U116** | 0.953 | 0.971 | 0.957 | 0.013 | 0.003 | 0.019 |
| **U121** | 0.971 | 0.981 | 0.980 | 0.010 | 0.011 | 0.009 |
| **A159** | 0.990 | 0.995 | 0.995 | 0.005 | 0.002 | 0.002 |
| **A166** | 0.992 | 0.993 | 0.991 | 0.002 | 0.003 | 0.004 |
| **U172** | 0.965 | 0.977 | 0.972 | 0.018 | 0.012 | 0.015 |
| **C174** | 0.878 | 0.924 | 0.908 | 0.044 | 0.043 | 0.059 |
| **U428** | 0.922 | 0.966 | 0.962 | 0.049 | 0.014 | 0.022 |
| **G436** | 0.906 | 0.946 | 0.941 | 0.029 | 0.018 | 0.020 |
| **C462** | 0.932 | 0.940 | 0.953 | 0.044 | 0.026 | 0.018 |
| **A468** | 0.930 | 0.949 | 0.940 | 0.018 | 0.012 | 0.034 |
| **A484** | 0.969 | 0.972 | 0.969 | 0.008 | 0.016 | 0.015 |
| **G509** | 0.946 | 0.961 | 0.948 | 0.027 | 0.028 | 0.038 |
| **A512** | 0.859 | 0.919 | 0.857 | 0.071 | 0.044 | 0.094 |
| **C517** | 0.991 | 0.988 | 0.989 | 0.001 | 0.008 | 0.005 |
| **A576** | 0.979 | 0.992 | 0.984 | 0.010 | 0.003 | 0.009 |
| **A590** | 0.973 | 0.989 | 0.988 | 0.013 | 0.000 | 0.003 |
| **G601** | 0.984 | 0.991 | 0.989 | 0.007 | 0.002 | 0.002 |
| **U627** | 0.955 | 0.951 | 0.947 | 0.017 | 0.031 | 0.009 |
| **G644** | 0.978 | 0.977 | 0.971 | 0.013 | 0.018 | 0.024 |
| **A668** | 0.974 | 0.973 | 0.974 | 0.005 | 0.010 | 0.010 |
| **G683** | 0.986 | 0.989 | 0.985 | 0.004 | 0.002 | 0.004 |
| ***C797*** | *0.847* | 0.937 | 0.925 | 0.027 | 0.015 | 0.015 |
| **U799** | 0.976 | 0.984 | 0.983 | 0.012 | 0.009 | 0.007 |
| ***G867*** | *0.754* | 0.935 | 0.939 | 0.022 | 0.015 | 0.006 |
| **A1031** | 0.974 | 0.982 | 0.984 | 0.021 | 0.014 | 0.011 |
| ***C1272*** | *0.609* | 0.726 | 0.747 | 0.024 | 0.066 | 0.037 |
| **U1288** | 0.958 | 0.961 | 0.962 | 0.028 | 0.036 | 0.033 |
| **U1326** | 0.969 | 0.971 | 0.964 | 0.005 | 0.014 | 0.010 |
| **G1328** | 0.960 | 0.965 | 0.963 | 0.014 | 0.012 | 0.007 |
| **A1383** | 0.977 | 0.979 | 0.932 | 0.009 | 0.016 | 0.059 |
| ***C1391*** | 0.968 | 0.923 | *0.823* | 0.015 | 0.043 | 0.082 |
| **U1442** | 0.946 | 0.981 | 0.970 | 0.039 | 0.009 | 0.019 |
| ***G1447*** | *0.000* | 0.248 | 0.158 | 0.000 | 0.131 | 0.113 |
| **G1490** | 0.999 | 0.999 | 0.999 | 0.000 | 0.000 | 0.000 |

| | | | | | | |
|---|---|---|---|---|---|---|
| **A1678** | 0.944 | 0.956 | 0.958 | 0.020 | 0.031 | 0.022 |
| **C1703** | 0.982 | 0.985 | 0.979 | 0.008 | 0.005 | 0.008 |
| **U1804** | 0.945 | 0.971 | 0.957 | 0.016 | 0.005 | 0.014 |

**TABLE S3: Average RMS scores at established 2'-O-Me sites in 28S rRNA from three donors.** Sites downregulated in SIPS cells in comparison with P/Q cells are highlighted in blue and italic and downregulated in P cells in comparison with Q/SIPS cells are highlighted in red and italic.

| | Average of HDF 76 & 161 & 85 | | | St. dev. HDF 76 & 161 & 85 | | |
|---|---|---|---|---|---|---|
| ***Established sites*** | **P** | **Q** | **SIPS** | **P** | **Q** | **SIPS** |
| **A389** | 0.973 | 0.981 | 0.973 | 0.012 | 0.007 | 0.012 |
| **A391** | 0.947 | 0.973 | 0.971 | 0.007 | 0.007 | 0.008 |
| ***G1303*** | *0.648* | 0.791 | 0.809 | 0.057 | 0.016 | 0.018 |
| **A1313** | 0.948 | 0.933 | 0.937 | 0.023 | 0.048 | 0.042 |
| ***C1327*** | 0.743 | 0.780 | *0.687* | 0.079 | 0.017 | 0.022 |
| **G1509** | 0.926 | 0.925 | 0.912 | 0.027 | 0.032 | 0.025 |
| **A1511** | 0.973 | 0.975 | 0.967 | 0.015 | 0.013 | 0.025 |
| **A1521** | 0.995 | 0.995 | 0.995 | 0.003 | 0.004 | 0.003 |
| **G1612** | 0.992 | 0.993 | 0.990 | 0.006 | 0.004 | 0.007 |
| **G1747** | 0.964 | 0.984 | 0.985 | 0.028 | 0.012 | 0.009 |
| **A1858** | 0.996 | 0.996 | 0.995 | 0.001 | 0.002 | 0.003 |
| **C1868** | 0.783 | 0.828 | 0.796 | 0.088 | 0.086 | 0.072 |
| **C2338** | 0.949 | 0.948 | 0.939 | 0.008 | 0.011 | 0.014 |
| **A2350** | 0.990 | 0.989 | 0.986 | 0.005 | 0.007 | 0.011 |
| **G2351** | 0.981 | 0.971 | 0.956 | 0.006 | 0.015 | 0.026 |
| **C2352** | 0.930 | 0.927 | 0.910 | 0.016 | 0.026 | 0.038 |
| **A2388** | 0.912 | 0.943 | 0.920 | 0.032 | 0.021 | 0.027 |
| **U2402** | 0.877 | 0.894 | 0.885 | 0.042 | 0.039 | 0.022 |
| **C2409** | 0.952 | 0.946 | 0.946 | 0.003 | 0.019 | 0.016 |
| **G2411** | 0.975 | 0.981 | 0.981 | 0.010 | 0.011 | 0.011 |
| **A2774** | 0.813 | 0.771 | 0.774 | 0.016 | 0.037 | 0.061 |
| **C2791** | 0.908 | 0.916 | 0.879 | 0.080 | 0.064 | 0.111 |
| **A2802** | 0.984 | 0.988 | 0.986 | 0.008 | 0.007 | 0.010 |
| **C2811** | 0.923 | 0.937 | 0.922 | 0.037 | 0.025 | 0.045 |
| **U2824** | 0.995 | 0.993 | 0.991 | 0.001 | 0.003 | 0.006 |
| **C2848** | 0.939 | 0.935 | 0.918 | 0.028 | 0.033 | 0.040 |
| **G2863** | 0.771 | 0.798 | 0.736 | 0.122 | 0.122 | 0.164 |
| ***C3680*** | 0.831 | 0.757 | *0.552* | 0.032 | 0.105 | 0.200 |
| **A3697** | 0.973 | 0.975 | 0.971 | 0.007 | 0.007 | 0.011 |
| **A3703** | 0.995 | 0.997 | 0.996 | 0.002 | 0.002 | 0.001 |
| ***G3723*** | *0.560* | 0.734 | 0.631 | 0.132 | 0.110 | 0.149 |
| **A3739** | 0.991 | 0.987 | 0.989 | 0.002 | 0.008 | 0.005 |
| **A3764** | 0.992 | 0.990 | 0.989 | 0.002 | 0.004 | 0.008 |
| **G3771** | 0.993 | 0.988 | 0.989 | 0.004 | 0.003 | 0.002 |

| | | | | | | |
|---|---|---|---|---|---|---|
| **C3787** | 0.987 | 0.986 | 0.985 | 0.008 | 0.010 | 0.010 |
| **U3797** | 0.980 | 0.983 | 0.968 | 0.022 | 0.018 | 0.038 |
| **A3804** | 0.982 | 0.982 | 0.979 | 0.004 | 0.002 | 0.004 |
| **A3809** | 0.970 | 0.974 | 0.966 | 0.008 | 0.007 | 0.008 |
| **C3820** | 0.982 | 0.981 | 0.980 | 0.008 | 0.009 | 0.007 |
| **A3846** | 0.917 | 0.910 | 0.914 | 0.011 | 0.008 | 0.023 |
| **C3848** | 0.880 | 0.902 | 0.918 | 0.072 | 0.065 | 0.040 |
| **C3866** | 0.948 | 0.959 | 0.945 | 0.011 | 0.006 | 0.014 |
| **G3878** | 0.980 | 0.978 | 0.980 | 0.006 | 0.011 | 0.008 |
| **U3904** | 0.972 | 0.967 | 0.973 | 0.016 | 0.024 | 0.015 |
| **G3923** | 0.864 | 0.865 | 0.860 | 0.083 | 0.105 | 0.102 |
| **G4020** | 0.839 | 0.876 | 0.871 | 0.094 | 0.098 | 0.090 |
| **C4032** | 0.971 | 0.965 | 0.967 | 0.012 | 0.020 | 0.021 |
| **G4166** | 0.983 | 0.985 | 0.981 | 0.004 | 0.001 | 0.002 |
| **U4197** | 0.997 | 0.996 | 0.997 | 0.001 | 0.001 | 0.000 |
| **G4198** | 0.976 | 0.962 | 0.965 | 0.009 | 0.023 | 0.020 |
| **U4276** | 0.987 | 0.986 | 0.984 | 0.003 | 0.003 | 0.003 |
| **G4340** | 0.977 | 0.984 | 0.980 | 0.005 | 0.007 | 0.004 |
| **G4362** | 0.959 | 0.963 | 0.963 | 0.017 | 0.009 | 0.005 |
| **C4426** | 0.965 | 0.972 | 0.963 | 0.005 | 0.003 | 0.006 |
| **G4464** | 0.994 | 0.996 | 0.996 | 0.003 | 0.003 | 0.001 |
| **U4468** | 0.974 | 0.974 | 0.977 | 0.013 | 0.023 | 0.013 |
| **G4469** | 0.973 | 0.972 | 0.971 | 0.009 | 0.013 | 0.008 |
| **A4493** | 0.993 | 0.994 | 0.993 | 0.004 | 0.005 | 0.003 |
| **C4506** | 0.996 | 0.996 | 0.996 | 0.000 | 0.000 | 0.001 |
| **A4541** | 0.956 | 0.977 | 0.982 | 0.018 | 0.009 | 0.005 |
| **A4560** | 0.940 | 0.962 | 0.964 | 0.020 | 0.014 | 0.016 |
| ***G4588*** | *0.638* | 0.779 | 0.719 | 0.063 | 0.042 | 0.043 |
| **U4590** | 0.863 | 0.864 | 0.848 | 0.019 | 0.010 | 0.007 |
| **G4593** | 0.977 | 0.957 | 0.948 | 0.004 | 0.019 | 0.025 |
| **G4607** | 0.706 | 0.742 | 0.758 | 0.062 | 0.058 | 0.024 |

**TABLE S4: Probes for northern blots.**

| Probe | snoRNA/ pre-rRNA | Oligo Sequence |
|---|---|---|
| hSNORD91B(HBII-296B) _G4588_probe_rev | snoRNA91A/B | AGTATCACACAGAAGTTGCATC |
| hSNORD91A(HBII-296A) _G4588_probe_rev | | GAACCACACAGAGATTGCAT |
| hSNORD87(HBII-276) _G3723_probe_rev | snoRNA87 | AGCTGGGTAAACGGCA |
| hSNORD21 _G1303_probe_rev | snoRNA21 | TGCCATCAGTCCCGTC |
| hSNORD28(U28) _ C1391_probe_rev | snoRNA28 | TGCCATCAGAACTCTAACAT |
| hSNORD127_G1447_probe_rev | snoRNA127 | GTTTAGAGGGACTGTTGTCCT |
| hSNORD88C_C3680_probe_rev | snoRNA88A/B/C | AGGTGTCAAAGGTCCTGG |
| hSNORD88B_C3680_probe_rev | | AGTGCTGGACATCACGG |
| hSNORD88A_C3680_probe_rev | | GTGCTGGACATCATGGAG |
| h5.8S-rRNA_ probe _rev | 5.8S rRNA | GTGTCCTGCAATTCACATT |
| hSNORD66_C1272_probe_rev | snoRNAD66 | GTTCCATCATGGTGTCTCAG |
| hSNORD98_G867_probe_rev | snoRNA98 | GTTCCACACTGCATTTCAG |
| hSNORD57_A99_probe_rev | snoRNA57 | GTCAGGCTCAGACAGTTCAT |
| OMK1874 | ITS2 | GGCAAGAGGAGGGCGGA |
| OMK1873 | ITS1 | GTCCGGGCTCCGTTAATGATC |

**TABLE S5: qPCR primers**

| Name | Primer type | Oligo Sequence |
|---|---|---|
| UniLoop | universal reverse primer | GTGCAGGGTCCGAGGT |
| snoR98 | Stem-loop_RT primer | GTTGGCTCTGGTGCAGGGTCCGAGGTATTCGCAC CAGAGCCAACGAGTTC |
| | Forward primer | GGGGGAGTTATGATGTGTGTAAATC |
| snoR28 | Stem-loop_RT primer | GTTGGCTCTGGTGCAGGGTCCGAGGTATTCGCAC CAGAGCCAACTGCCAT |
| | Forward primer | TGGGTCAGATGATTTGAATTGATAAG |
| snoR66 | Stem-loop_RT primer | GTTGGCTCTGGTGCAGGGTCCGAGGTATTCGCAC CAGAGCCAAC TTCCTC |
| | Forward primer | GTGTGTTTCCTCTGATGACTTCC |
| snoR127 | Stem-loop_RT primer | GTTGGCTCTGGTGCAGGGTCCGAGGTATTCGCAC CAGAGCCAACTGGCAA |
| | Forward primer | GTTGGCAACTGTGATGAAAGAT |
| snoR21 | Stem-loop_RT primer | GTTGGCTCTGGTGCAGGGTCCGAGGTATTCGCAC CAGAGCCAACGCTGCC |
| | Forward primer | TGGGCTGAATGATGATATCCCA |
| snoR88C | Stem-loop_RT primer | GTTGGCTCTGGTGCAGGGTCCGAGGTATTCGCAC CAGAGCCAACCTGGGG |
| | Forward primer | GTTTCTGGGGCTCCCATGAT |
| snoR87 | Stem-loop_RT primer | GTTGGCTCTGGTGCAGGGTCCGAGGTATTCGCAC CAGAGCCAACTCTCAG |
| | Forward primer | TGGGGACAATGATGACTTAAATTACTTTT |
| snoR91B | Stem-loop_RT primer | GTTGGCTCTGGTGCAGGGTCCGAGGTATTCGCAC CAGAGCCAACAAAAGCC |
| | Forward primer | TGGGGAAGAGCCAATGATGTTTTTAT |
| 5S | Forward primer | CATACCACCCTGAACGCG |
| | Reverse primer | CTACAGCACCCGGTATTCCC |
| 5.8S | Forward primer | ACTCTTAGCGGTGGATCA |
| | Reverse primer | ATCAATGTGTCCTGCAATTC |
| GAPD | Forward primer | CGACCACTTTGTCAAGCTCA |
| | Reverse primer | TGTGAGGAGGGGAGA TTCAG |
